# Application of HPMCAS-coated Ctx(Ile^21^)-Ha peptide microparticles as a potential use to prevent systemic infection caused by *Salmonella* Enteritidis in poultry

**DOI:** 10.1101/2021.03.19.436179

**Authors:** Cesar Augusto Roque Borda, Mauro de Mesquita Souza Saraiva, Daniel F. M. Monte, Lucas Bocchini Rodrigues Alves, Adriana Maria de Almeida, Taísa Santiago Ferreira, Túlio Spina de Lima, Valdinete Pereira Benevides, Julia Memrava Cabrera, Andréia Bagliotti Meneguin, Marlus Chorilli, Angelo Berchieri Junior, Eduardo Festozo Vicente

## Abstract

The transmission of *Salmonella* Enteritidis (SE) in poultry is most often by the fecal-oral route, which can be attributed to the population density. Consequently, the pathogen triggers stress response and virulence factors deploying it to survive in hosts. Therefore, this study proposed to evaluate HPMCAS-coated microparticles containing the Ctx(Ile^21^)-Ha antimicrobial peptide against SE in laying hens chicks’ infection model to determine whether Ctx(Ile^21^)-Ha-utilization confers a benefit in the intestinal lumen, as well as whether limits systemic infection. Importantly, while assessing whether AMP utilization confers reduction of SE in liver, it was noted that there was statistical significance between groups A (control, no Ctx(Ile^21^)-Ha peptide) and B (2.5 mg of Ctx(Ile^21^)-Ha/kg) at 2 dpi, potentially indicating the Ctx(Ile^21^)-Ha effectiveness in the first stage of infection by SE. Remarkably, it was also detected a statistical significance (*p* -value <0.0001) with lower counts of SE (∼ 0 CFU) in livers at 5, 7, and 14 dpi, regardless of Ctx(Ile^21^)-Ha dosage (2.5 mg or 5 mg/kg - group C). By using Chi-square test, the AMP effect on SE fecal excretion was evaluated. In this regard, it was noticed statistical significance (*p* < 0.05) among groups B and C in comparison with control group A, since those groups had lower bacterial excretion along 21 days. In summary, the role of HPMCAS-Ctx(Ile^21^)-Ha peptide microcapsules against *S*. Enteritidis in laying hen chicks infection model was unraveled, providing a satisfactory results against this pathogen.

## Introduction

*Salmonella enterica* subsp. *enterica* serovar Enteritidis is one of the leading cause of foodborne diseases posing global concerns to one health and economy ^1–3^. In this concern, several efforts have been made to reduce the contamination and spread of *Salmonella* Enteritidis along the poultry production chain, since this serovar is most often associated with poultry products ^4,5^. Besides that, the overuse of antimicrobial agents in animal husbandry, have contributed to the emergence of virulent and multidrug resistant (MDR) strains among poultry products, which represents a critical public health issue, once it could have implications on human health ^2,6,7^.

The increasing spread of virulent and MDR strains have imposed the poultry production sector to decrease the use of antimicrobial agents and simultaneously to find alternative solutions to mitigate such pathogen ^8,9^. Among these mitigation strategies, the use of non-conventional drugs such as antimicrobial peptides (AMPs) has been recognized to have anti-*Salmonella* effect, not only for MDR strains, but especially against high virulent *Salmonella*. These promising AMPs are molecules that are able to modulate the immune response, which protect hosts against invasive infections ^8,9^.

Antimicrobial peptides triggers destabilization of bacterial cell membrane preventing their growth, which could possibly inhibit the lethality of *Salmonella* spp. Consequently, the antibacterial mechanisms of AMPs have become a research hotspot ^3^. Interestingly, gut inflammation provides a growth advantage for *Salmonella*, contributing to becoming this pathogen more harmful ^10,11^. Indeed, the virulence package plays a crucial role in invasive non-typhoidal *Salmonella* (NTS) infections, favoring their growth and survival in hosts ^10,11^. Therefore, this work proposed to evaluate the HPMCAS-coated microparticles of Ctx(Ile^21^)-Ha peptide against *S*. Enteritidis in laying hen chicks infection model to determine whether Ctx(Ile^21^)-Ha-utilization confers a benefit in the intestinal lumen, as well as whether limits systemic infection.

## Materials and methods

### Chemical agents

Hypromellose Acetate Succinate (HPMCAS, AQOAT® - Grade AS-LF; Shin-Etsu Chemical Co., Ltd), Fmoc-aminoacids, Rink Amide resin, *N,N′-*Diisopropylcarbodiimide (DIC; CAS No. 693-13-0), Hydroxybenzotriazole (HOBt; PubChem SID 57651485), Hexahydropyridine (CAS No. 110-89-4), Trifluoroacetic acid (TFA; CAS Number: 76-05-1), Triisopropylsilane (TIS; #233781) and Acetonitrile (ACN; #34851) in High performance liquid chromatography (HPLC)/analytical grade and Phosphate-buffered saline (PBS; #P5493) were purchased from Sigma-Aldrich.

Dimethylformamide (DMF; Neon Comercial #01114), dichloromethane (DCM; Anidrol Products Laboratories #PAP.A-1986), sodium alginate (#3913.10.00) and aluminum chloride (#2827.32.00) were obtained from Êxodo Científica, Brilliant Green agar (BGA; K25-610009) culture medium and selenite cystine broth (SCB; K25-610150) were purchased KASVI, Sodium Biselenite (#2030) was acquired from INLAB, MacConkey agar (MC; #CM0007B) and BD Difco™ LB Broth (LB; # DF0402-07-0) was obtained from Fisher Scientific.

### Ctx(Ile^21^)-Ha antimicrobial peptide assembly

Ctx(Ile^21^)-Ha (MW = 2,289.72 g mol^-1^) was synthesized in solid phase manually and characterized as previously described ^12^. Briefly, all amino acids and resin were protected with fluorenylmethoxycarbonyl α-aminic protecting group (Fmoc) ^13^. Rink amide resin (degree of substitution = 0.6 mmol g^-1^) was employed as the solid support for the peptide synthesis, presented by the following sequence GWLDVAKKIGKAAFSVAKSFI-NH ^14^. Fmoc-amino acids were coupled for 2 h with DIC/HOBt (0,6 equivalents) previously dissolved on ultrasound in 1:1 DMF/DCM. Afterwards, the protector was removed with 2:8 Hexahydropyridine/DMF to able the coupling of the next amino acid.

Once the peptide construction was concluded, the resin was separated from the peptide, using a cleavage solution of 95% TFA, 2.5% TIS, and 2.5% water and stirred for 2 h at room temperature. The solution was precipitated three times with cold diethyl ether and both phases were separated with a Pasteur pipette, manually, centrifuged, and drying in the desiccator with silica beads. For the extraction of the peptide, were used HPLC mobile phases to solubilize the crude peptide, which was lyophilizated after the process. Ctx(Ile^21^)-Ha identification was performed by HPLC (Shimadzu with membrane degasser DGU-20A5R, UV detector SPD-20A, column oven CTO-20A, automatic sampler SIL-10AF, fraction collector FRC-10A and LC-20AT dual-pump, C18 column) and characterization by Mass Spectrometry (Bruker Amazon, Brazil), using a mobile phases proportion of A (0.045% TFA/H2O) and B (0.036% TFA/ACN), 1:1, v/v, at 220/280 nm wavelength detection.

### Obtaining of HPMCAS-coated microparticles

#### Ionic gelation^15^

2% sodium alginate aqueous solution was homogenized with Ctx(Ile^21^)-Ha peptide for 4 h, until complete dissolution. Then, using a pump and a syringe, the alginate was cross-linked dropwise in aluminum chloride and they were placed in a drying oven for 6 h at 40°C. Crosslinking solutions had the final concentrations of peptide: B = 0.2 g L^-1^ and C = 0.4 g L^-1^.

#### Enteric coating^16^

The microparticles obtained were placed in a fluidized bed at 40°C, with a peristaltic pump speed of 0.4 mL min^-1^, system vibration at 100%, and a 0.25 L min^-1^ blower. The enteric coating solution was prepared with HPMCAS, ammonium hydroxide, triethylcitrate and water (1:2.5:0.25:6.25, w/v).

### Bacterial strain

The *Salmonella* Enteritidis strain P125109 (accession number AM933172.1) used in this study was isolated from an outbreak of human food poisoning in the United Kingdom ^17,18^. The stock culture of SE strain kept at −80°C was aerobically cultured at 37°C in Lysogeny Broth (LB) for 18 h. After incubation, an aliquot (100 µL) was serially diluted (1:10) in Phosphate Buffered Saline pH 7.4 (PBS) and inoculated onto LB agar plates. Thereafter, the inoculum of SE strain was maintained approximately at 108 CFU cell suspension before oral infection.

### Ethical statements and *in vivo* assays

*In vivo* experiments were performed according to the Ethical Principles on Animal Experimentation (CEUA) of the National Council for the Control of Animal Experimentation (CONCEA). The protocol was approved by the Ethical Committee on Animal Experimentation from the School of Agriculture and Veterinarian Sciences (FCAV) at August, 25^th^ 2020 (Protocol number: 2588/20). Experimental assays were carried out in the Department of Pathology, Theriogenology, and One Health at São Paulo State University (FCAV/UNESP).

To determine whether AMPs’ utilization confers protection against *Salmonella* Enteritidis, one hundred and twenty laying hen chicks were obtained from a commercial hatchery. Upon arrival, the presence of *Salmonella* spp. was investigated by using cloacal swab, as well as drag swab of transport cardboard boxes ^19^. Samples were dispensed in sterile Selenite Broth containing 0.04% of Novobiocin (SN – Becton Dickinson, Maryland, USA) and incubated at 37°C for 24h. Afterwards, they were inoculated onto MacConkey and Brilliant Green agar, subsequently being incubated at 37°C for 24h.

Next, randomly distributed 40 laying hen chicks per group, including control group A (no Ctx(Ile^21^)-Ha; n=40), B (2.5 mg of Ctx(Ile^21^)-Ha/kg; n=40), and C (5 mg of Ctx(Ile^21^)-Ha/kg; n=40) were performed. Laying hen chicks were orally infected with 0.2 mL (10^8^ CFU/mL) of *S*. Enteritidis.

Chicks were euthanized 2, 5, 7, 14, 21 days after infection. Ceacal contents were collected for enumeration of SE, whereas liver and spleen were collected to evaluate the systemic infection, also by using bacterial counts. *S*. Enteritidis counts were determined by plating serial ten-fold dilutions onto BG agar, supplemented with nalidixic acid (100 µg/mL) and spectinomycin (100 µg/mL).

### Fecal excretion

Fifteen chicks were randomly selected and subjected to cloacal swabs, which were performed twice a week during 21 days period. The collected cloacal swabs were dispensed in 2 mL of SN Broth and incubated for 24 h at 37°C. Subsequently, they were streaked onto BG agar (BGA – Oxoid, UK) containing 100 µg/mL of nalidixic acid and 100 µg/mL of spectinomycin being incubated for 24 hours at 37°C. Presumptive colonies of *Salmonella* spp. were confirmed positives by tests performed in slide agglutination using *Salmonella* O Poly Antisera (anti-O – Bio-Rad, USA).

### Statistical analysis

Fisher’s exact test was used to compare *Salmonella* excretion on the feces between the groups (P< 0.05). While the logarithmically transformed values for bacterial counts from cecal content, liver, and spleen, were submitted to two-way ANOVA followed by Bonferroni multiple comparison test (P < 0.05). All statistical analyses were performed using the software GraphPad Prism, version 8.2.1.

## Results

### Ctx(Ile^21^)-Ha synthesis and microparticles results

HPLC and LC/MS analyzes and microparticles characterization and physicochemical stability were previously published elsewhere ^12^.

Stable and light yellow microparticles were obtained (shown in Figure 1), total mass of 20.05 and 20.01 g of non-coated microparticles, and 24.51 and 23.96 g of HPMCAS-coated microparticles, for B and C, respectively. The microparticles obtained had an average size of 2 mm, which served not to be very different from common poultry food and was not visibly rejected or preferred for them. The mean final concentration of each coated capsule was 3.1 and 6.3 µg Ctx(Ile^21^)-Ha per mg of microparticles for B and C, respectively.

**Figure 1.**
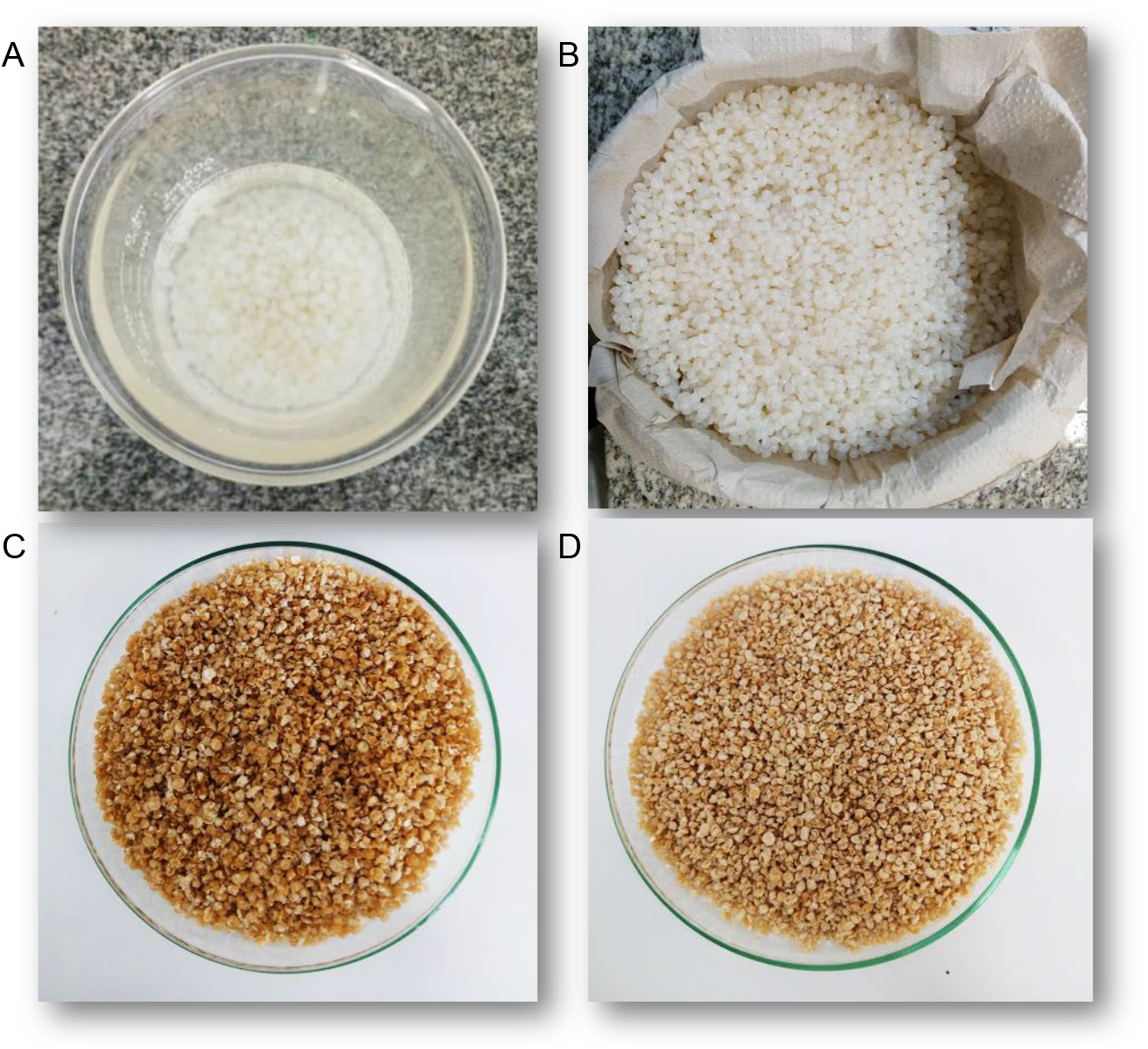
**A**. Microcapsules in solution after ionic gelation. **B**. Pre-dried isolated non-encapsulated microparticles. **C**. B-microparticles coated with HPMCAS of Ctx(Ile^21^)-Ha peptide (3.1 µg/mg). **D**. C-microparticles coated with HPMCAS of Ctx(Ile^21^)-Ha peptide (6.3 µg/mg).

### Effect of Ctx(Ile^21^)-Ha antimicrobial peptide on *Salmonella* Enteritidis cecal colonization

While assessing whether Ctx(Ile^21^)-Ha AMP utilization confers protection against *S*. Enteritidis, it was evidenced statistical difference in ceacal content only in 5 days post infection (dpi), where A (control group) had higher counts of SE when compared with groups B (*p-*value = 0.0392) and C (*p-*value = 0.0056) according to Bonferroni’s multiple comparisons (BMC) test as summarized in Figure 2. It was also noted higher counts of SE in ceacal content of the group A in comparison with B and C in 14 dpi, but this difference did not reach statistical significance (Figure 2).

**Figure 2.**
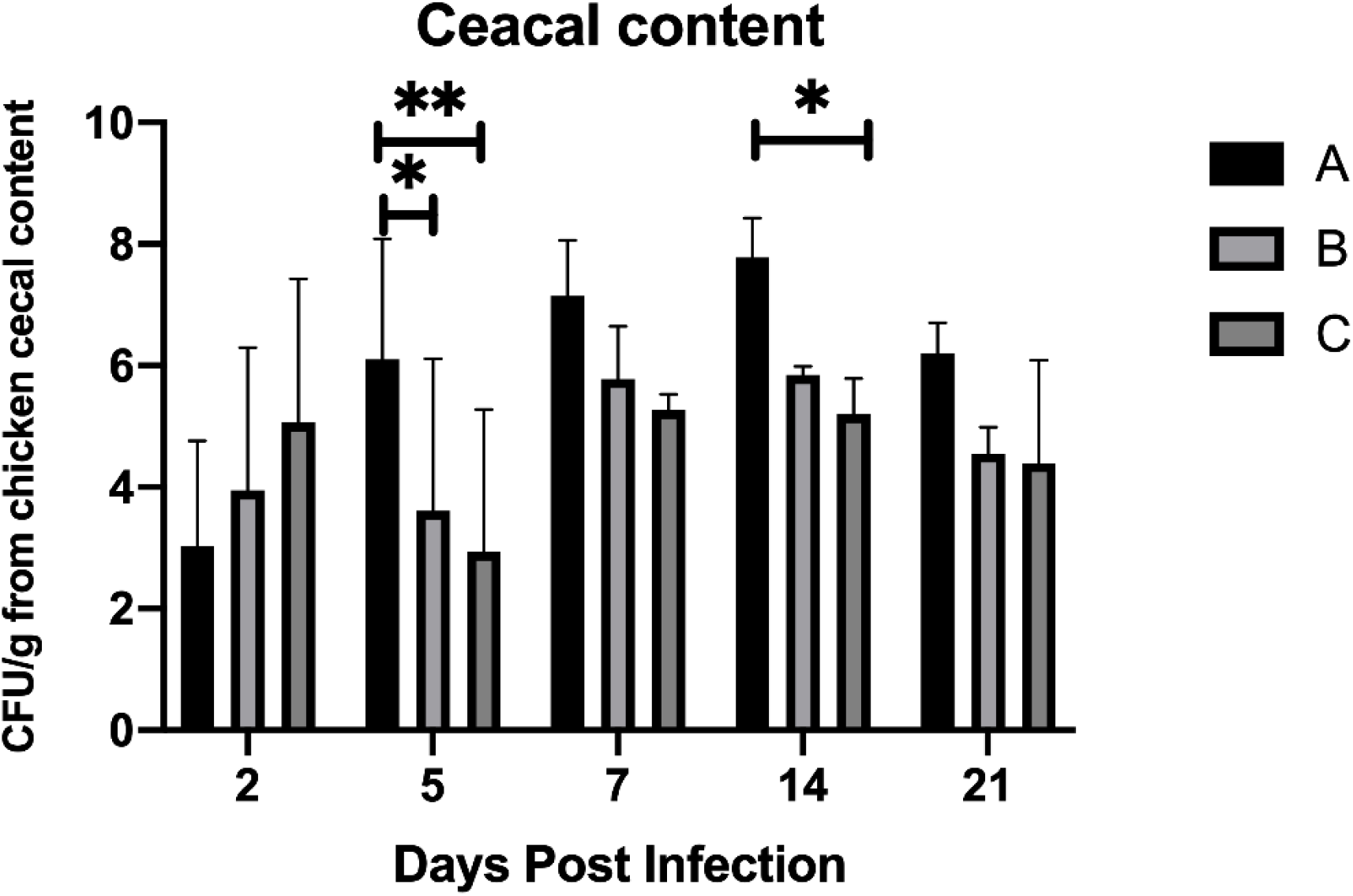
Effect of antimicrobial peptides on *Salmonella* Enteritidis cecal colonization.

### Contribution of the Ctx(Ile^21^)-Ha to avoid *Salmonella* Enteritidis systemic infection

Similarly, for systemic infection analysis, it was seen statistical significance of group C in 5 (*p*-value = 0.0095) and 14 (*p*-value = 0.0103) dpi compared to the control group (A), which demonstrated that 5 mg of Ctx(Ile^21^)-Ha AMP can reduce the bacterial counts in spleen (Figure 3). In addition, two-way ANOVA test showed significant difference as bacterial count between the treatments of each group (Interaction *p-*value = 0.0262, Row Factor *p-*value = 0.0005, Column Factor *p-*value = 0.1772).

**Figure 3.**
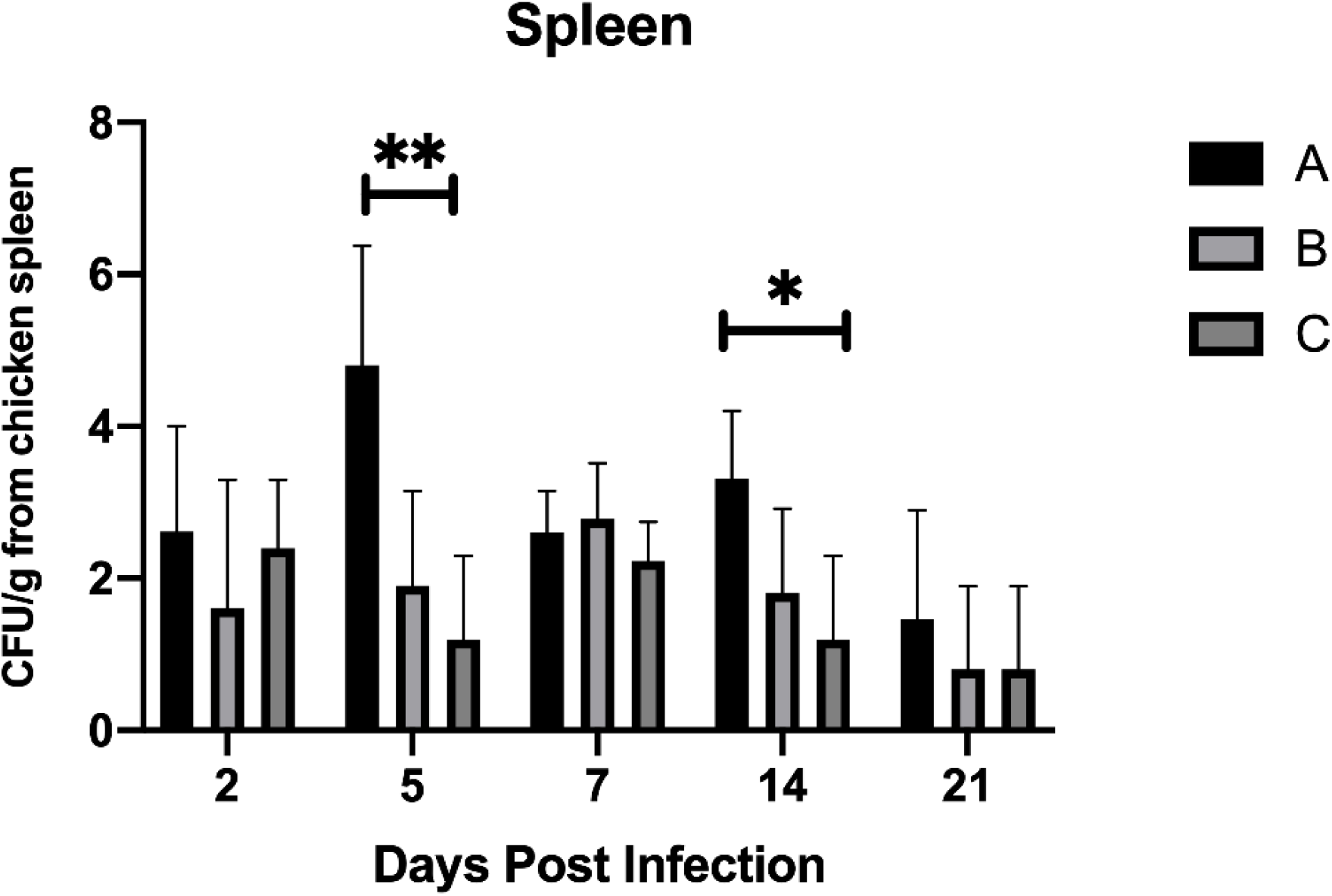
Effect of antimicrobial peptides on *Salmonella* Enteritidis spleen infection.

Importantly, while assessing whether AMPs utilization confers reduction of *S*. Enteritidis in liver, it was noted that there was statistical significance (A vs. B, *p-*value <0.0001; B vs. C, *p-*value = 0.0021) between groups B and A at 2 dpi, potentially indicating the Ctx(Ile^21^)-Ha effectiveness in the first stage of infection by *S*. Enteritidis (Figure 4). Remarkably, it was also evidenced a statistical significance (*p*-value <0.0001) with lower counts of SE (∼ 0 CFU) in livers at 5, 7, and 14 dpi, regardless of Ctx(Ile^21^)-Ha dosage (2.5 mg or 5 mg), as shown in Figure 4.

**Figure 4.**
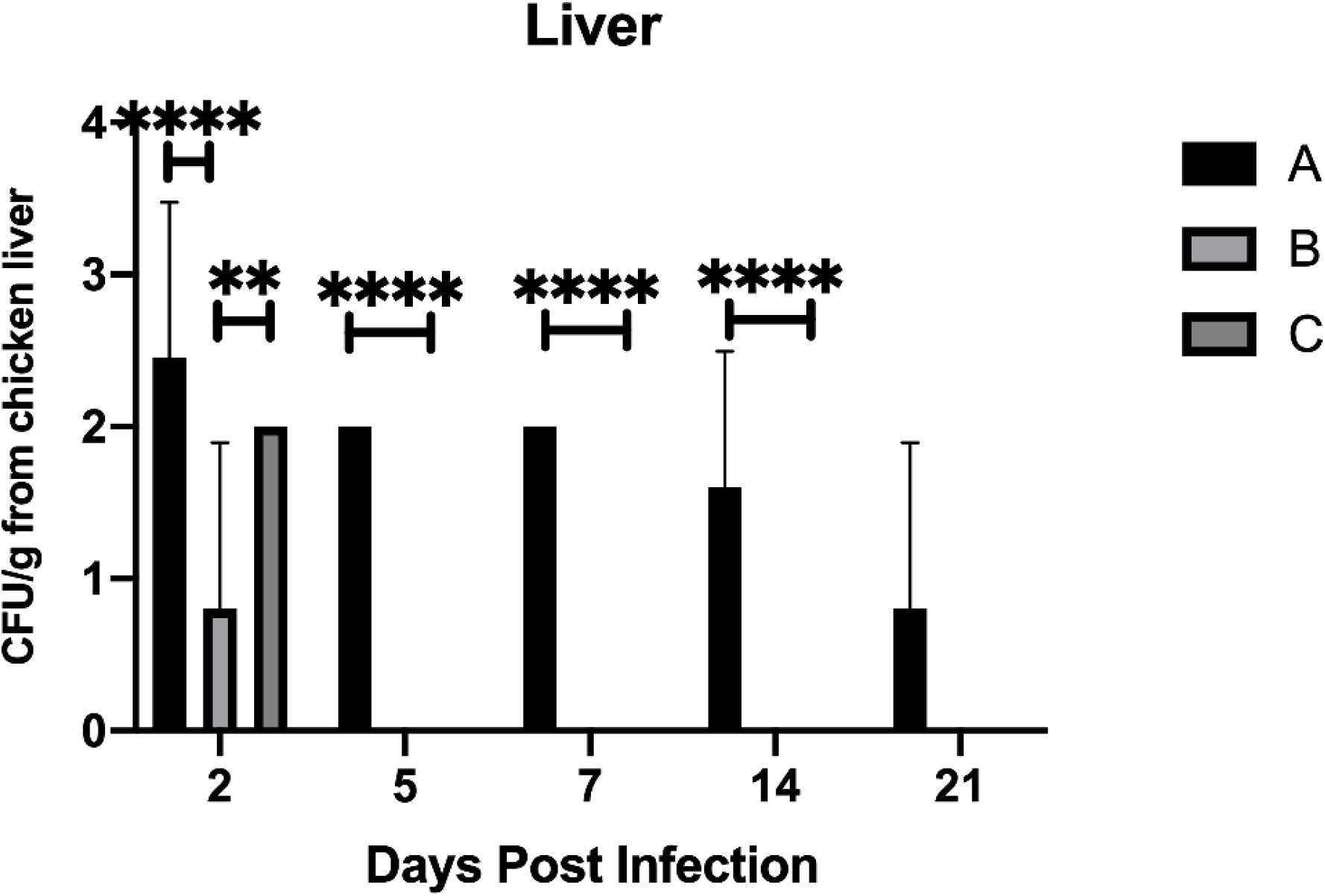
Effect of antimicrobial peptides on *Salmonella* Enteritidis liver infection.

Additionally, two-way ANOVA showed a significant difference in all groups (Interaction *p-*value = 0.0062, Row Factor *p-*value <0.0001, Column Factor *p-*value <0.0001), which confirms this anti-Systemic Infection potential against SE.

### Fecal excretion

By using Chi-square test, the effect of Ctx(Ile^21^)-Ha antimicrobial peptide on *S*. Enteritidis fecal excretion was evaluated. In this regard, it was noticed statistical significance (*p* < 0.05) among groups B and C in comparison with control group A, since those groups had lower bacterial excretion along 21 days, as shown in Figure 5.

**Figure 5.**
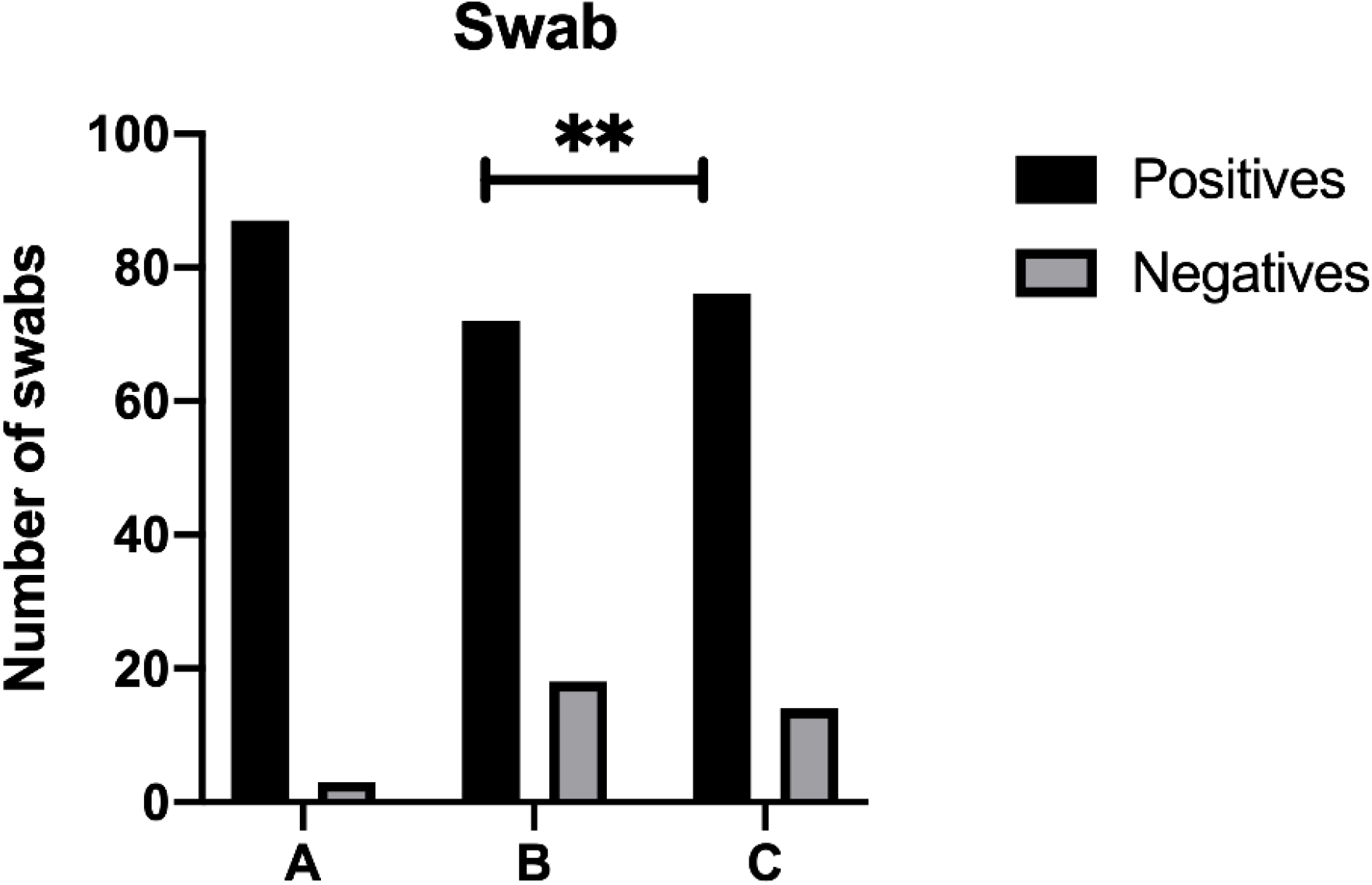
Evaluation of the effect of antimicrobial peptides on *Salmonella* Enteritidis fecal excretion during 21 days.

## Discussion

Peptides and proteins are biomolecules that require special attention when used as oral drugs, because they must travel throughout the gastrointestinal tract (GIT). Due to this, they can be easily denatured, deactivated or hydrolyzed by the presence of proteases or an acid environment (low pH) ^20^. It has been shown that in the gastric tract (pH ∼ 1-3), the main problem with peptides would be related to the presence of pepsin (10 – 15 % hydrolysis). In this way, due to constant shear, its stability would be completely affected. In addition, the enzymes present in the intestine (trypsin, chymotrypsin, aminopeptidase, etc.) would be responsible for the total breakdown of peptide bonds ^20,21^. The challenges in the transport of the AMPs through the GIT made this treatment not entirely efficient ^22^.

HPMCAS is a pH-dependent biopolymer, resistant to pH-acid, so the Ctx(Ile^21^)-Ha peptide was protected during its passage through the stomach. However, in the intestinal pH, HPMCAS is totally soluble and the coating entirely dissolves ^16^. Carboxyl radicals in solution help the temporary inactivation of GIT enzymes, which can chelate metals used as cofactor i.e., Mg^+2^ and Ca^+2 23,24^. Most of the enzymatic factors were solved using HPMCAS and alginate, because they have several carboxyl radicals in their chemical structure (Figure 6).

**Figure 6.**
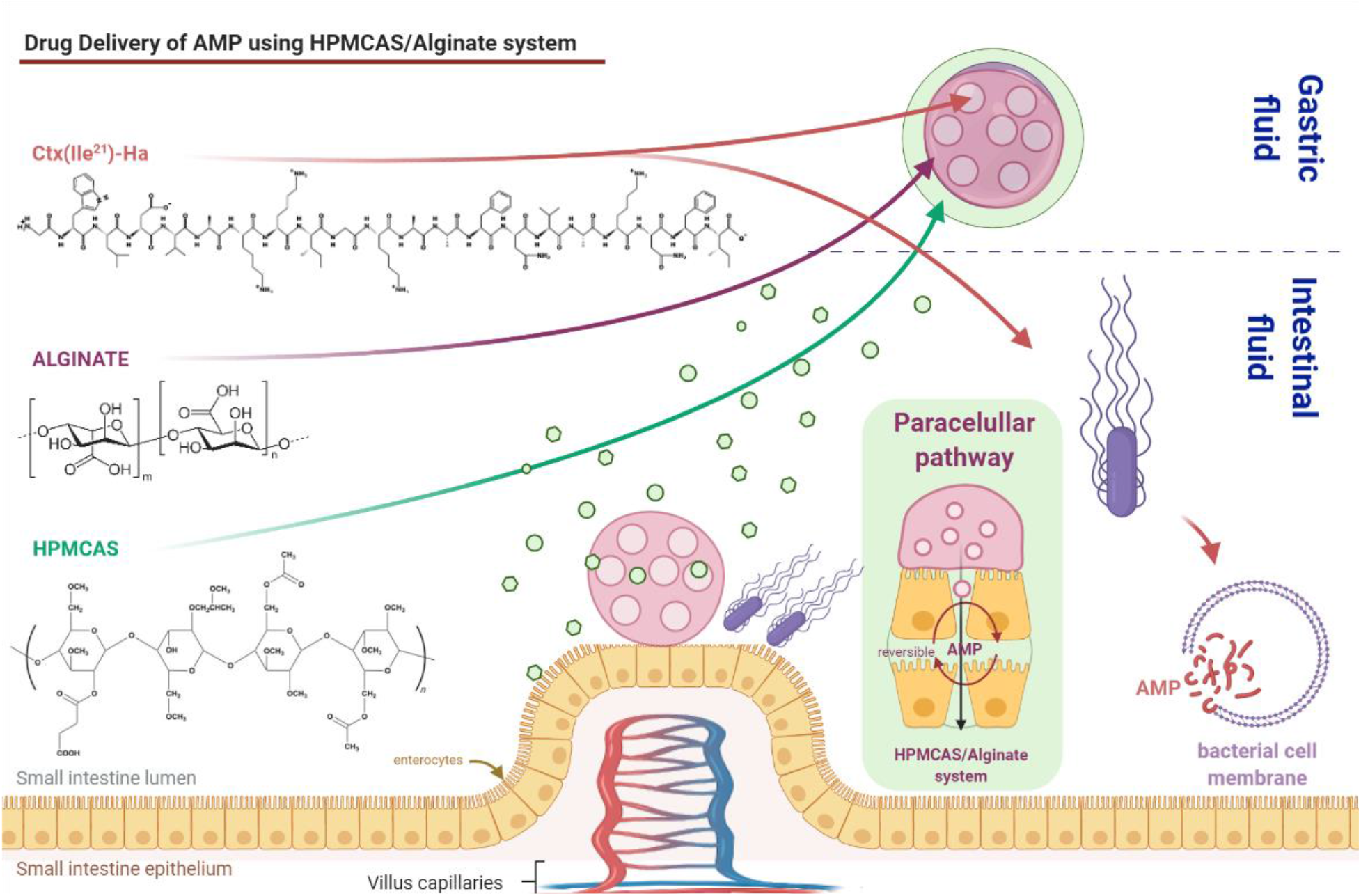
Brief explanation of the absorption pathway and antibacterial activity of the antimicrobial peptide Ctx(Ile^21^)-Ha.

Intestinal absorption of AMP (Figure 6), due to its size, low permeability and the physical barriers of the mucosa, could be an obstacle ^25,26^. However, it has been shown that the use of biopolymers could allow these molecules to pass through these compartments without difficulty. In this way, use of polymers with mucoadhesive and mucopenetration capacity are necessary ^27^. In a previous study, the absorption of insulin, which has a molecular weight even higher than that of the Ctx(Ile^21^)-Ha peptide, was achieved using the same ionic gelation techniques, with a combined pectin/retrograded starch system, which successfully crossed the paracellular pathway and was demonstrated that polymeric microparticles are capable of opening tightness of tight junctions facilitating the transport of peptides ^28^.

In this investigation, the system was favored by the mucoadhesive ^29^, mucopenetration ability ^30^, and biocompatibility of the alginate ^31^, increasing the bioavailability of the peptide ^32^. Furthermore, previous studies indicate that HPMCAS was able to improve permeation through monolayers of human colon adenocarcinoma cells (Caco-2) and dialysis membranes ^33^. This polymer was also able to inhibit the crystallization of drugs (mainly by succinyl groups) and to keep them stable in a solid phase dispersion ^34,35^. This evidence corroborates the results obtained regarding the Ctx(Ile^21^)-Ha peptide mechanism of action in the liver and could explain the high rate of bacterial elimination.

AMPs have several mechanisms of interaction with the cell membrane lipopolysaccharides that manage to destabilize the bacteria, reaching a high biological potential, even with MDR bacteria ^36^. Ctx(Ile^21^)-Ha belongs to the ceratotoxins’ family, originally isolated from the skin of a Brazilian Cerrado frog ^37^. Previous studies indicated potential antimicrobial activity against SE and *Salmonella enterica* subps. *enterica* serovar Typhimurium (ST) and other MDR bacteria ^12^.

Other AMPs were studied to inhibit the replication of *Salmonella* sp., such as cathelicidin-BF, which demonstrated an anti-ST effect in murine with a Minimum Inhibitory Concentration (MIC) of 1.1 mol L^-1 22^. However, no values were shown for anti-SE, being more efficient results compared to anti-ST (64 μg mL^-1^) of Ctx(Ile^21^)-Ha ^12^.

Ctx(Ile^21^)-Ha has also anti-SE MIC values of 4 mol L^-1^, better than indolicidin AMP (8,4 mol L^-1^). Therefore, its potential is maintained. Furthermore, human cathelicidin LL-37, a largely studied peptide with potential activity, showed low anti-ST activity (28 mol L^-1^), but did not show anti-SE activity. Ctx(Ile^21^)-Ha demonstrated better anti-SE results when compared to conventional drugs, such as Chlortetracycline (36 mol L^-1^) and Neomycin (13 mol L^-1^) ^38^. However, such studies remain scarce. In this regard, this study proposes a novel product with combined techniques from food chemistry and biotechnology and pharmaceutical fields.

Notably, *S*. Enteritidis remains one of the most important pathogens associated with poultry products ^4,5^. In this regard, several efforts made by poultry industry have been reduced the cross-contamination along the food chain. Consequently, these mitigation strategies have benefited the human health by avoiding the food poisoning caused by this high priority pathogen. Despite this, the development of new efficient antimicrobials is still scarce ^39^. In this concern, antimicrobial peptides have been recognized as a viable alternative against pathogenic bacteria ^40^. However, in some circumstances, virulent lineages are able to colonize the ceca environment and may also spread through liver, spleen, and heart, causing systemic infection ^41^. Presumably, this statement corroborates our findings, since chicks with no AMP treatment (group A) substantially presented more bacterial counts than treated chicks (groups B and C). Therefore, this study could provide the potential use of Ctx(Ile^21^)-Ha antimicrobial peptide to reduce *S*. Enteritidis counts in chicken infection model.

One of the most important aspects related to the successful application of this HPMCAS against *S*. Enteritidis, is most likely regarding to the high mechanical resistance ^42^ and insolubility in gastric pH of the molecule, which could guarantee the fully release of Ctx(Ile^21^)-Ha peptide ^15^.

Besides, increasing evidences have demonstrated that treatment with AMPs would generate a better immune response during the first 5 days of life ^43,44^. The use of AMPs, such as bacteriocins in poultry production, could prevent reinfection ^45^, through factors that advance the immune response ^46^. Therefore, our results corroborate the decrease in bacterial load with a significant difference in the intestine in the first days after infection (5 dpi), as shown in Figure 2.

Care in food handling and animal production should be a priority, without using drugs excessively, since Salmonella as well as other bacteria are prone to the acquisition of bacterial resistance and consequently produce a reinfection with more serious effects ^47,48^. However, the presented results of fecal excretion showed a relevant significant difference since the complete elimination of SE was achieved from a large number of chickens, without the presence of reinfection. This arises a different vision of the use of these biopharmaceutical additives that could replace conventional drugs to control the systemic infection of SE and other subspecies, with minimal risk of bacterial resistance.

## Conclusions

Collectively, these data demonstrated that the use of formulated antimicrobial peptides, particularly the Ctx(Ile^21^)-Ha, could be a promising alternative against systemic infections caused by *S*. Enteritidis, deserving to be more explored against other *Salmonella enterica* serovars. To the best of our knowledge, this is the first report of an Ctx(Ile^21^)-Ha peptide that displayed satisfactory results against *S*. Enteritidis in laying hen chicks’ infection model. This outcome might be useful at animal husbandry as a plausible alternative with anti-*Salmonella* effect.

## Data Availability statement

The data that support the findings of this study are openly available in Mendeley Data at http://doi.org/10.17632/cgkt7pxj2n.1

## Acknowledgments

This work was developed with the support of São Paulo Research Foundation/FAPESP (Process number 2016/00446-7) and master scholarship (Process number 2018/25707-3). We thank the technical assistants of Laboratory of Chemistry and Biochemistry from São Paulo State University (Unesp), School of Sciences and Engineering, Tupã; University of Araraquara which made available the use of the fluidized-bed equipment and Shin-Etsu company for gently donate and provide the HPMCAS coating for the experiments. Finally, the research group “Peptides: Synthesis, Optimization and Applied Studies - PeSEAp”. This work is a chapter of master’s thesis and is also part of the national patent protected throughout the Brazilian territory by the National Institute of Intellectual Property (INPI BR1020200220489).

